# A ubiquitous mobile genetic element disarms a bacterial antagonist of the gut microbiota

**DOI:** 10.1101/2023.08.25.553775

**Authors:** Madeline L. Sheahan, Michael J. Coyne, Katia Flores, Leonor Garcia-Bayona, Maria Chatzidaki-Livanis, Anitha Sundararajan, Andrea Q. Holst, Blanca Barquera, Laurie E. Comstock

## Abstract

DNA transfer is ubiquitous in the gut microbiota, especially among species of Bacteroidales. *In silico* analyses have revealed hundreds of mobile genetic elements shared between these species, yet little is known about the phenotypes they encode, their effects on fitness, or pleiotropic consequences for the recipient’s genome. Here, we show that acquisition of a ubiquitous integrative and conjugative element encoding an antagonistic system shuts down the native contact-dependent antagonistic system of *Bacteroides fragilis*. Despite inactivating the native antagonism system, mobile element acquisition increases fitness of the *B. fragilis* transconjugant over its progenitor by arming it with a new weapon. This DNA transfer causes the strain to change allegiances so that it no longer targets ecosystem members containing the same element yet is armed for communal defense.

## Main text

Bacteroidales is an order of bacteria that includes numerous species that colonize the human gut at high density (*1, 2*), many likely persisting for decades (*3*). Two distinct types of contact-dependent interactions are pervasive among gut Bacteroidales; conjugal DNA transfer, and antagonism mediated by Type VI secretion systems (T6SS). T6SSs are contractile nanomachines where a pointed tube loaded with toxin effectors is injected into neighboring cells. Of the numerous species of this order, *Bacteroides fragilis* is one of the most antagonistic, producing several diffusible secreted anti-bacterial toxins that thwart intra-species competitors (*4–7*), as well as a broad targeting T6SS produced only by this species (*8*).

The *B. fragilis* specific T6SS, known as genetic architecture 3 (GA3), is present in ∼ 75% of strains analyzed (*9*) and robustly antagonizes diverse Bacteroidales species *in vitro* and *in vivo* (*10–12*). Two other T6SS loci are present in gut Bacteroidales species, GA1 and GA2, and are each present on large integrative and conjugative elements (ICE) that readily transfer between Bacteroidales species in the human gut (*8, 9*). Of the numerous mobile genetic elements (MGE) of the gut Bacteroidales (*9, 13*), the GA1 and GA2 ICE are among the few with a conserved architecture that are commonly spread to multiple species within individuals of industrialized populations (*9*). Strong antagonism (1-3 log killing) has not been demonstrated for either the GA1 or GA2 T6SSs despite the presence of genes encoding potent toxins in these loci (*8, 14, 15*). There are few transcriptomic or phenotypic studies analyzing the fate of Bacteroidales MGEs once transferred to a new bacterial host (*15a*,*16, 29*), or their impact on the recipient’s genome, transcriptome and proteome (*17*).

*Bacteroides* are saccharolytic bacteria, with most species preferentially utilizing dietary plant polysaccharides that reach the colon largely untouched by human enzymes (*18, 19*). *B. fragilis*, however, has a relatively limited nutritional niche, utilizing only a few plant polysaccharides (*20*), and preferentially utilizing host mucin glycans (*21*). When dietary polysaccharides are scarce, other *Bacteroides* species can switch to consume host mucin glycans (*21*), likely placing them in range of the broadly targeting GA3 T6SS of *B. fragilis*.

Prior analyses revealed that the *B. fragilis* GA3 T6SS fires constitutively *in vitro* (*10*) and without apparent directionality (*22*). We began by determining if constitutive *in vitro* firing is a general property of *B. fragilis* GA3 T6SSs. T6SS firing in broth grown cultures is assayed by the presence of the tube protein, Hcp, in the culture supernatant of strains grown to stationary phase. We assayed for the presence of the main structural Hcp protein in cells and culture supernatants of a panel of *B. fragilis* strains with distinct effector and immunity genes in the two GA3 T6SS locus variable regions (Fig. 1A). Stark differences were observed in the amount of Hcp present in supernatants (Fig. 1B); most notable are strains 1284, 2_1_56FAA, and S36_L11 that synthesize but do not fire the T6SS under these *in vitro* conditions. The ∼25 kb GA3 T6SS loci of these three strains are very similar to those of strains that constitutively fire (fig. S1), some with only a few nucleotide variations, suggesting that a factor(s) encoded outside the GA3 locus abrogates firing in these strains. A prior study showed that 18% of *B. fragilis* strains analyzed contain both GA3 and GA1 T6SS loci (*9*). Genomic analysis of the panel of *B. fragilis* strains showed that five of the GA3-containing *B. fragilis* strains also contain a GA1 T6SS ICE in their genomes (fig. S2), including all three that do not fire. These data suggest that, in these strains, a product of the GA1 ICE may inhibit firing of the GA3 T6SS.

**Fig. 1.**
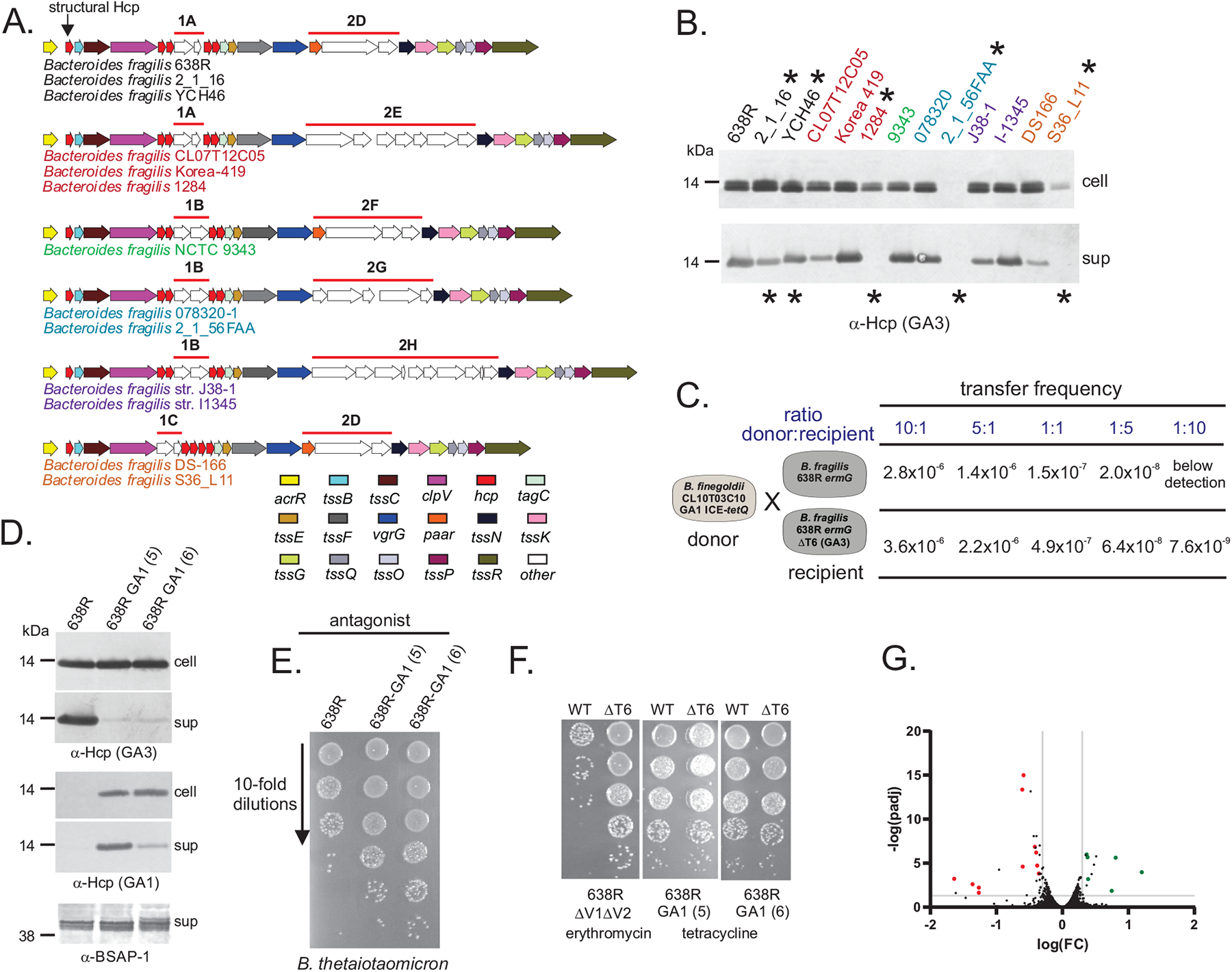
Acquisition of a GA1 ICE shuts down firing and antagonism of the *B. fragilis* GA3 T6SS. **(A)** Gene maps of the GA3 T6SS loci of 13 *B. fragilis* strains. Genes are color coded as shown below. Strain names are colored the same if they have the same variable region 1 and 2 (V1, V2), which contain genes encoding effector and immunity proteins. These 13 strains contained 3 different V1 (1A-1C) and five different V2 regions (2D-2H). **(B)** Western immunoblot analysis of the presence of the GA3 T6SS needle protein in the cell fraction or the supernatant. The same color coding used in panel A is used for these strains. Strains designated with an * have a GA1 T6SS ICE in their genomes. **(C)** Transfer frequencies of the GA1 ICE from donor strain *finegoldii* CL09T03C10 to either *B. fragilis* 638R or 638RΔT6, deleted for genes necessary for firing the GA3 T6SS. The *B. fragilis* strains have the *ermG* gene at a null site and the *B. finegoldii* GA1 ICE contains the t*etQ* gene at a null site to allow for selection of transconjugants. Five different ratios of donor:recipient were assayed for transfer frequency (fig. S3). **(D)** Western immunoblot analysis of the cellular fraction and supernatants of 638R and 638R-GA1 transconjugants 5 and 6. **(E)** Antagonism assays showing that the two 638R-GA1 transconjugants are defective in killing *B. thetaiotaomicron* VPI-5482 compared to the WT strain. **(F)** Antagonism assays with the transconjugant strains as the target strains to analyze their protection. 638RΔT6 is unable to fire and antagonize and 638RΔV1ΔV2 is deleted for both of the GA3 variable regions and is unable to protect itself (*10*). Following co-culture to allow antagonism, 638RΔV1ΔV2 was selected based on integration of the *ermG* gene into the chromosome, and 638R-GA1 transconjugants are selected based on tetracycline resistance encoded in the GA1 ICE. **(G)** Volcano plot showing the differentially expressed genes of 638R-GA1 transconjugant 6 compared to WT 638R. The red or green dots identify genes that are downregulated or upregulated, respectively, in both 638R-GA1 transconjugants 5 and 6 (transconjugant 5 volcano plot shown in fig. S4). Full RNASeq data from broth grown bacteria provided in Dataset S1.

The fact that a substantial number (∼18%) of GA3-containing *B. fragilis* strains contain GA1 ICE suggests that the GA3 T6SS may not be effective in preventing conjugal DNA transfer from targeted (antagonized) strains, unlike what has been shown in some bacterial species (*23, 24*). To study the ability of GA1 ICE to prevent GA3 firing and the contribution of the GA3 T6SS in preventing conjugal transfer, we quantified the transfer frequency of a GA1 ICE from donor strain *Bacteroides finegoldii* CL09T03C10 (BfineCL09) to a *B. fragilis* strain that constitutively fires its GA3 T6SS (638R), and to an isogenic strain that cannot fire this weapon (638R ΔT6). We found only a 1.5-3.2-fold reduction in transfer of the GA1 ICE due to GA3 antagonism when five different ratios of donor/target (BfineCL09) to recipient/antagonist (638R or 638RΔT6SS) were analyzed (Fig. 1C, fig. S3). When the recipient/antagonist were in equal abundance or at 5- and 10-fold excess to the donor/target, there was substantial GA3 T6-mediated killing of BfineCL09 (fig. S3), yet conjugal transfer was only modestly reduced, possibly due to delayed death of the donor strain following injection of toxins.

The GA1 ICE integrates into a 6-bp consensus site (*9*), allowing for its insertion throughout recipient genomes. Analysis of two resulting transconjugants revealed that the GA1 ICE of 638R-GA1 transconjugant 5 inserted between BF638R_4113-4114, and in 638R-GA1 transconjugant 6, between BF638R_1192-1193, both far from the GA3 T6SS locus which comprises genes BF638R1970-1994. Western blot analysis of the cells and supernatants of these 638R-GA1 transconjugants confirmed that acquisition of a GA1 ICE inhibits firing of the GA3 T6SS (Fig. 1D). Concurrent with this lack of GA3 firing, both strains lost the ability to antagonize (Fig. 1E), confirming that acquisition of a GA1 ICE prevents GA3 firing and antagonism. Assays with 638R as antagonist and each 638R-GA1 transconjugant as target, showed that both transconjugant strains were able to protect themselves from GA3 attacks (Fig. 1F), suggesting that these transconjugants produce GA3 immunity proteins *in vitro*. Transcriptomic analysis of WT 638R and 638R-GA1-5 and 638R-GA1-6 grown *in vitro* showed remarkably little genome-wide transcriptional differences due to acquisition of the GA1 ICE (Fig. 1G, fig S4, dataset S1) with no significant differences in expression of any of the GA3 T6SS locus genes (dataset S1). Therefore, lack of GA3 T6SS firing *in vitro* in 638R-GA1 is not due to decreased transcription of GA3 genes. An operon of genes (Bf638R_1955-1952) was significantly downregulated due to acquisition of the GA1 ICE, yet when deleted from the WT 638R strain, the GA3 T6SS still fired (fig. S5), showing that downregulation of these genes is does not inhibit firing. Transcriptomic analyses also revealed that the GA1 T6SS locus genes are expressed (dataset S1) and the GA1 system fires in these transconjugants (Fig. 1D), despite their inability to antagonize *in vitro* (Fig. 1E).

To identify genes of the GA1 ICE responsible for preventing firing of the GA3 T6SS, we deleted three large regions of the GA1 ICE in 638R-GA1-6, which collectively encompass most of the GA1 ICE except the left side which contains *tra* genes and other genes commonly found on Bacteroidales ICE (Fig. 2a). Western blot analysis of the cell lysates and supernatants of these mutants showed that deletions 1 and 2 did not affect GA3 firing, but deletion 3, which removed the GA1 T6SS region and some flanking genes, restored GA3 firing (Fig. 2C) and the strain’s ability to antagonize (Fig. 2D). A smaller deletion of this region (deletion 4) that removed only genes of the GA1 T6SS locus, similarly, restored GA3 firing (Fig. 2C) and antagonism (Fig. 2D), showing that *in vitro* firing of the GA3 T6SS is inhibited by genes of the GA1 T6SS locus. To determine if GA3 firing can be restored by smaller deletions in the GA1 T6SS locus, we made seven deletions spanning this locus (Fig. 2B) and found only partial restoration by deletion of *tssG-R* and *tssC-clpV* (Fig. 2E), most of which encode proteins of the baseplate and transmembrane complex. A surprising finding is that the GA1 Hcp needle is fired in these deletion mutants, suggesting that the system may be able to utilize the heterologous GA3 T6SS machinery.

**Fig. 2.**
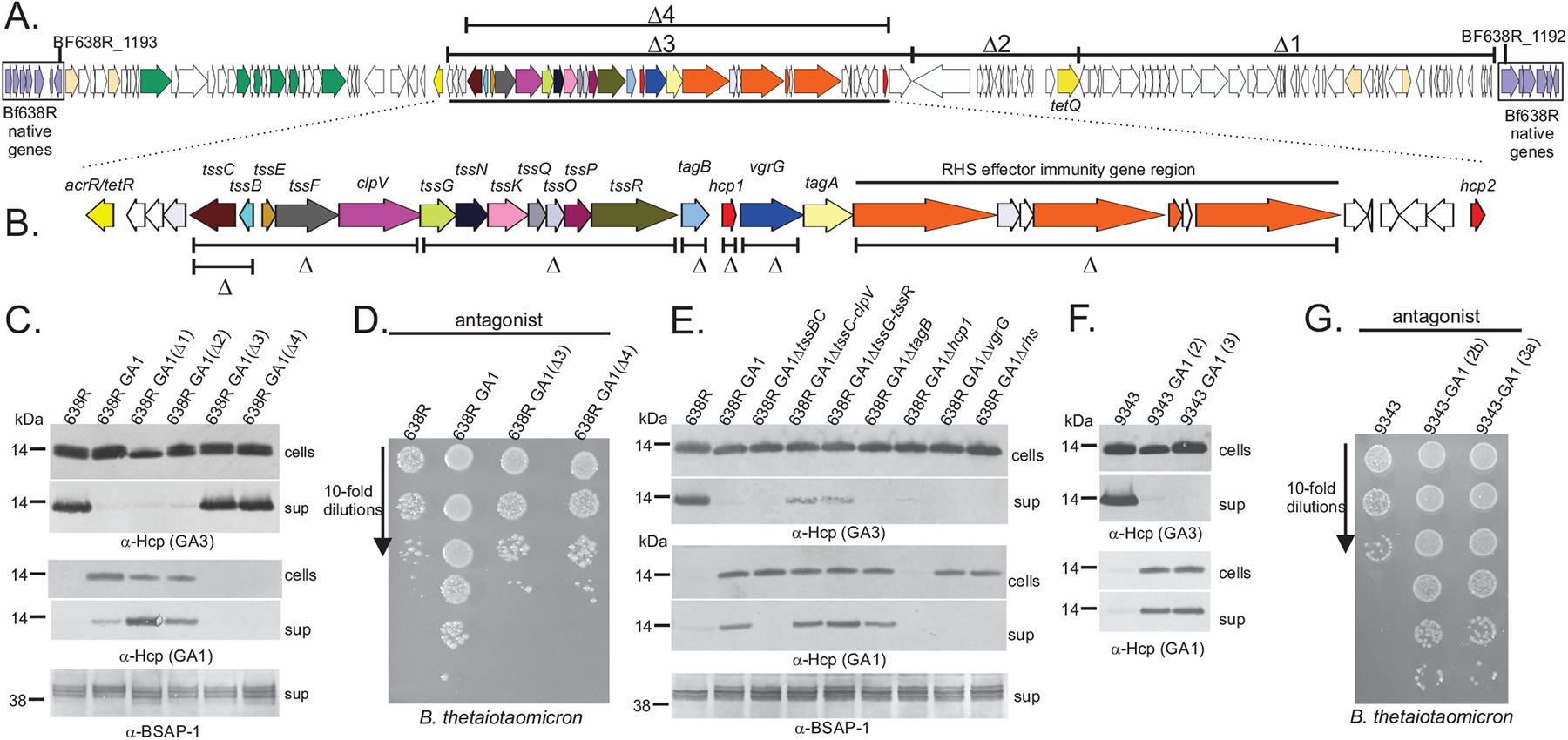
Identification of GA1 ICE region responsible for quelling GA3 firing. **(A)** ORF map of the GA1 ICE of *B. finegoldii* CL09T03C10 showing the region of the *B. fragilis* chromosome where the ICE inserted in 638R-GA1 transconjugant 6. Genes of the GA1 T6SS are color coded as the GA1 ICE shown in fig. S2. Genes on the left side of the ICE involved in conjugation processes are colored green. *tetQ* added to the ICE to allow selection of transconjugants is highlighted gold. Large deletions made within this region in 638R-GA1 are indicated. The region comprising the GA1 T6SS locus is underlined. **(B)** Deletions made throughout the GA1 T6SS locus in 638R-GA1. **(C)** Western immunoblots of cell lysates and supernatants of WT and 638R-GA1 and large deletion mutants indicated in panel A, probed with antisera to the GA3 and GA1 Hcp proteins, or an antiserum to BSAP-1 which is a secreted protein of 638R used as a loading control. **(D)** Antagonism assay using the strains listed on the top as the antagonist and *B. thetaiotaomicron* used as the target strain. The plates contain cefoxitin and are selective for *B. thetaiotaomicron*. **(E)** Western immunoblot performed similar in a manner similar to panel C except with the mutants shown in panel B. **(F)** Western immunoblot analysis of the cellular fraction and supernatants of *B. fragilis* 9343 and 9343-GA1 transconjugants 2b and 3a. **(G)** Antagonism assays showing that the two 9343-GA1 transconjugants are defective in killing *B. thetaiotaomicron* compared to WT 9343.

To show that GA1 inhibition of GA3 firing is a general property, we transferred the GA1 ICE from BfineCL09 to *B. fragilis* 9343 (fig. S6), which has different GA3 effector and immunity genes in the two GA3 locus variable regions compared to 638R (Fig. 1B). The resulting transconjugants (9343-GA1) were also unable to fire the GA3 T6SS (Fig. 2F) and to antagonize (Fig. 2G), demonstrating a general inhibitory function by GA1 ICE on GA3 T6SS firing.

The *in vitro* results above show that the 638R-GA1 strain no longer carries the energetic cost of firing the GA3 T6SS yet is still able to protect itself from GA3 toxicity when targeted by the ancestral strain. Based solely on these GA3-based properties, the 638R-GA1 strain may be more fit in a competition assay. To determine if GA1 ICE acquisition by 638R affects its fitness *in vivo*, we performed a gnotobiotic mouse competitive gut colonization assay, competing 638R against 638R-GA1. Mice were gavaged with a 1:1 mixture of the strains, and bacteria were quantified from feces two-and four-weeks post-gavage. By two weeks, 638R-GA1 strain outcompeted the 638R stain in all six mice (Fig. 3A), with 638R below the detection level in four of the six mice, and by 4 weeks, no 638R were detected in any mice (Fig. 3A). Therefore, in this model, the acquisition of the GA1 ICE provides a competitive advantage to GA3-containing *B. fragilis* strains. When we assayed for the presence of the GA1 and GA3 Hcp proteins in the fecal material, the results were unexpected and drastically differed from the *in vitro* results. In all samples in which strain 638R was below the level of detection, no GA3 Hcp was detected in the feces (Fig. 3B), suggesting that 638R-GA1 does not produce GA3 Hcp *in vivo* as it does *in vitro*. In the two fecal samples from day 14 where 638R was present at 42% and 8% of the total bacteria, there was detectable GA3 Hcp (Fig. 3B), showing that 638R produces GA3 Hcp *in vivo*, but that acquisition of the GA1 ICE prevents its *in vivo* synthesis. As conclusive proof that 638R-GA1 does not produce GA3 Hcp *in vivo*, we mono-colonized gnotobiotic mice with either 638R or 638R-GA1, where they colonized at similar levels of approximately 2 x 10^10^ – 3 x 10^10^ and confirmed that synthesis of GA3 Hcp is abrogated *in vivo* in 638R-GA1 (Fig. 3C). Passage of 638R-GA1 *in vitro* after recovery from the mono-colonized mouse showed restoration of GA3 Hcp synthesis (fig. S7), demonstrating that lack of production *in vivo* was not due to selection of GA3 mutants. These collective data show that in this *in vivo* model, the GA1 ICE not only inhibits GA3 T6SS firing as it does *in vitro*, but completely abrogates GA3 Hcp synthesis.

**Fig. 3.**
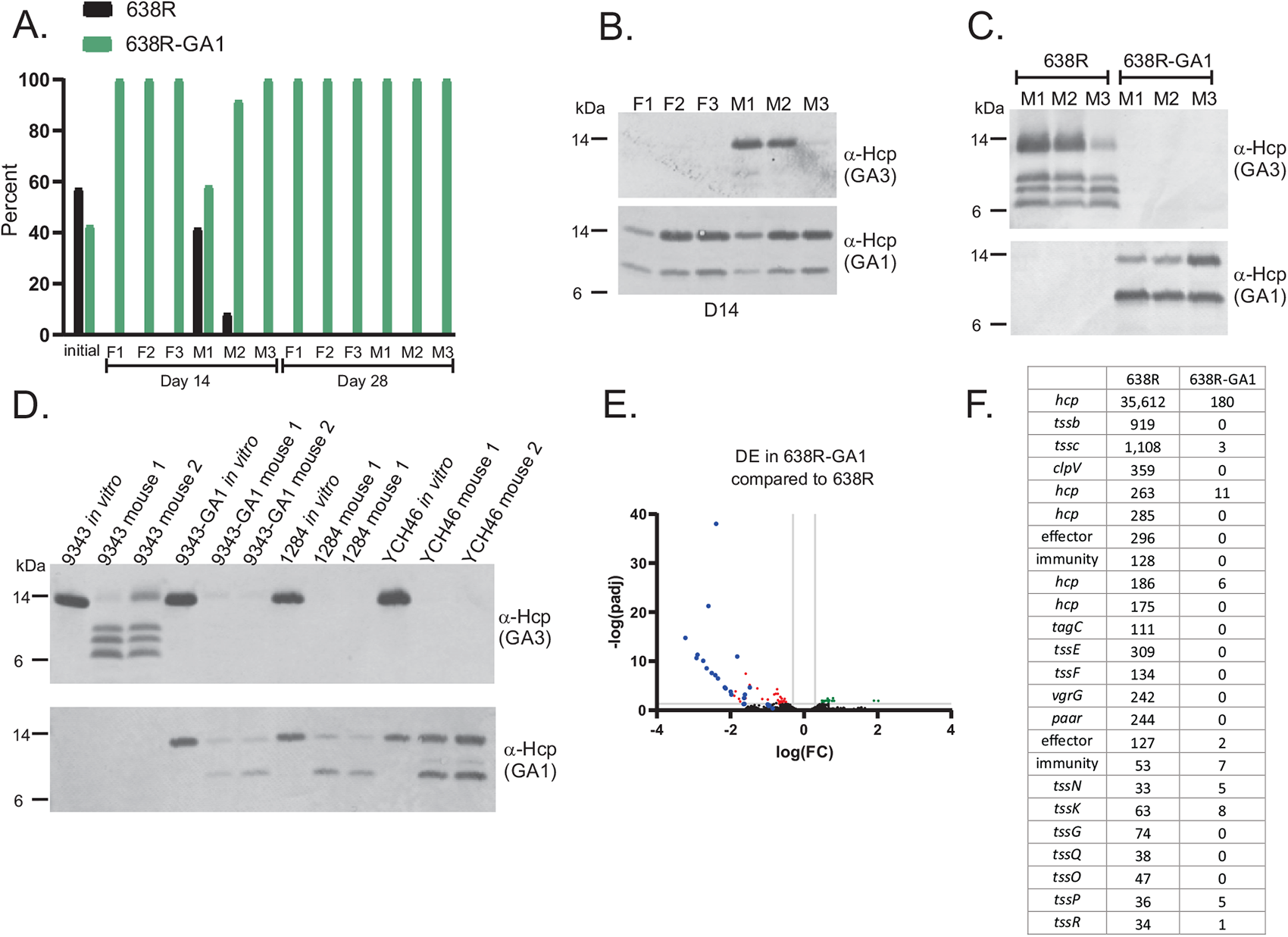
GA1 effects on the 638R GA3 T6SS in the mammalian gut. (**A)** Competitive colonization assay in gnotobiotic mice using three female (F) and three male (M) mice showing the percentage of the two strains in the initial inoculum and then in the feces at day 14 and day 28 post-gavage. (**B)** Western immunoblot analysis of day 14 fecal samples from mice from the competition experiment in panel A. **(C)** Western immunoblot analysis of fecal samples from three mice mono-colonized with either 638R or 638R-GA1 after 7 days of colonization. **(D)** Western immunoblot analysis of cells from broth-grown bacteria or from fecal samples from mice mono-colonized with three additional *B. fragilis* strains containing a GA3 T6SS and a GA1 ICE. **(E)** Volcano plot showing differential expression of genes of 638R-GA1 compared to 638R from feces of mice monocolonized with each strain. Blue dots indicate genes of the GA3 T6SS. Red dots indicate other significantly downregulated genes and green dots indicate significantly upregulated genes (significance defined as at least 2.0-fold and adjusted P value (DESeq2) and FDR (edgeR) both were less than or equal to 0.05). **(F)** Transcript per kilobase million (TPK) values for the GA3 T6SS genes from feces of mice monocolonized with 638R or 638R-GA1.

To determine if *in vivo* inhibition of GA3 synthesis is a general property of *B. fragilis* strains that have both a GA3 locus and GA1 ICE, we mono-colonized mice with 9343, 9343-GA1, or two *B. fragilis* strains, 1284 and YCH46, that naturally contain both a GA3 locus and a GA1 ICE (Fig. 1A, fig. S2). *B. fragilis* YCH46 was one of the two strains with a GA1 ICE that was able to fire the GA3 T6SS *in vitro* (Fig. 1B). We found that none of the strains that naturally contain a GA1 ICE, nor those that received the GA1 ICE by *in vitro* transfer, produce the GA3 Hcp protein *in vivo* (Fig. 3D), confirming that the GA1 ICE encodes a function that shuts down synthesis of GA3 Hcp *in vivo*. Transcriptomic analysis of fecal samples from mice mono-colonized with 638R or 638R-GA1 revealed robust transcription of the GA3 genes from 638R, but a near compete abrogation of transcription of GA3 T6SS genes in 638R-GA1 (dataset S2, Fig. 3E, 3F).

In our initial analyses of the three distinct genetic architectures of T6SS loci in gut Bacteroidales (*8*), we noted that each contain an adjacent gene encoding a transcriptional repressor of the AcrR/TetR family (*25*). In the BfineCL09 GA1 ICE, this gene is four genes downstream of *tssC* (Fig. 2A). To determine if AcrR_GA1_ is involved in the repression of GA3 transcription in 638R-GA1, we deleted *acrR*_GA1_ from this strain and mono-colonized mice with this mutant. Western blot analysis of feces following one-week mono-colonization showed restoration of synthesis of GA3 Hcp (Fig. 3A). To broaden the relevance of this discovery, we deleted *acrR*_GA1_ from 9343-GA1, and from the natural GA1-GA3 containing strain Bf1284 and found that the synthesis of GA3 Hcp is restored *in vivo* when *acrR*_GA1_ is deleted in all these GA3, GA1 ICE strains (Fig 3A), whether created in the lab or isolated from nature. Transcriptional analysis of 638R-GA1Δ*acrR*_GA1_ from mono-colonized mouse feces revealed that the genes that are significantly downregulated *in vivo* due to acquisition of the GA1 ICE are restored when *acrR*_GA1_ is deleted (dataset S2, Fig. 3B), such that comparison of the *in vivo* transcriptome of 638R with that of 638R-GA1Δ*acrR*_GA1_ showed only two differentially expressed 638R genes (dataset S2, Fig. 4C). An interesting finding is that deletion of *acrR*_GA1_ decreased transcription of one of the three operons (*hcp*-*rhs*) of the GA1 T6SS locus (dataset S2), revealing that this transcriptional repressor indirectly enhances rather than represses GA1 T6SS genes.

**Fig. 4.**
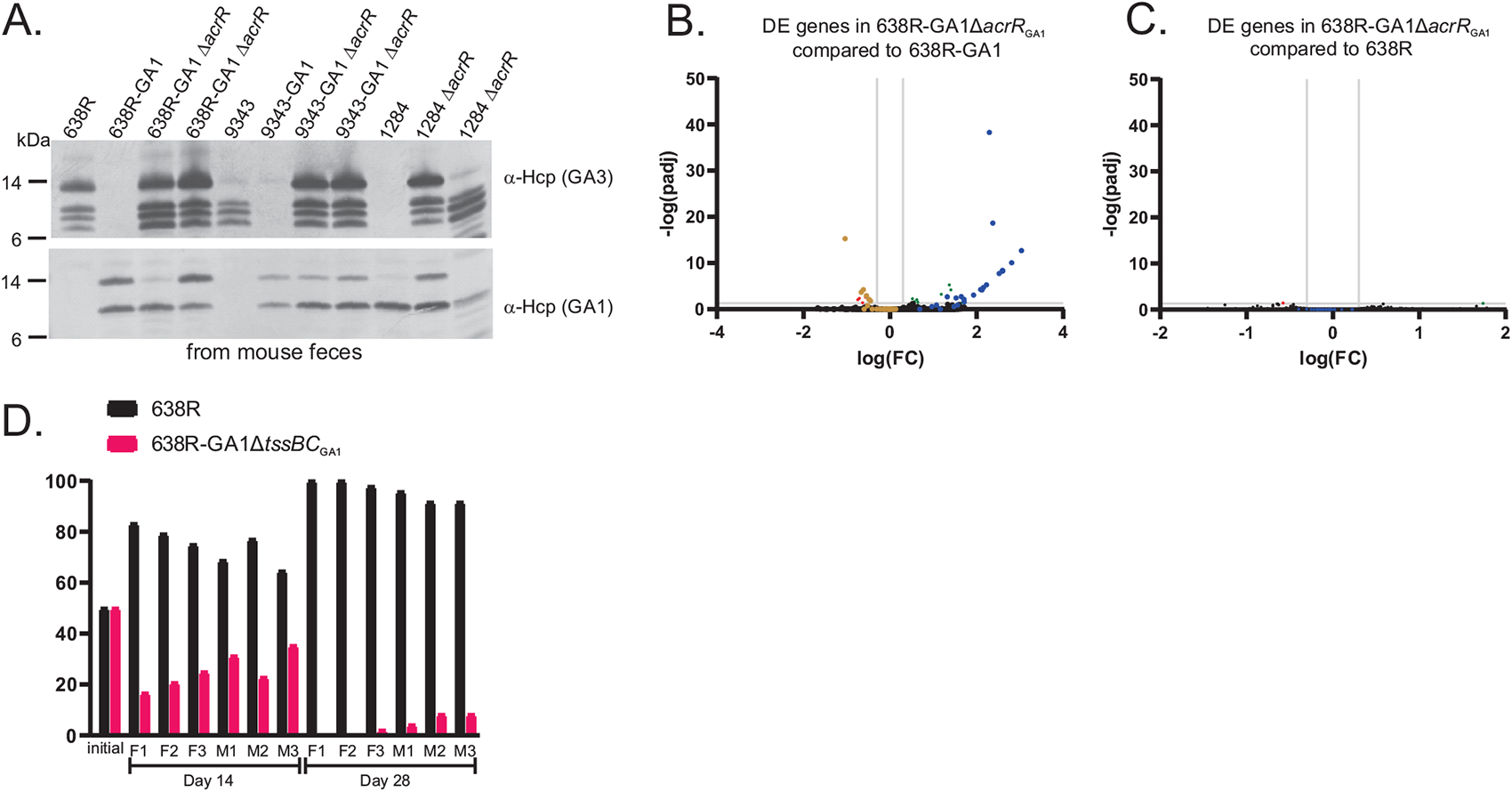
Analyses of regulation and competition *in vivo*. **(A)** Western immunoblot analysis of Hcp synthesis in WT *B. fragilis* strains, GA3-GA1 *B. fragilis* strains, and *acrR*_GA1_ deletion mutants from feces of mono-colonized mice. **(B)** Volcano plot showing differential expression of genes of 638R-GA1Δ*acrR*_GA1_ compared to 638R-GA1 from feces of mice monocolonized with each strain. Blue dots indicate genes of the GA3 T6SS locus. Brown dots indicate genes of the GA1 T6SS locus (significance defined as at least ≥2 -fold and adjusted *p*-value (DESeq2) and FDR (edgeR) both were less than or equal to 0.05). **(C)** Volcano plot showing differential expression of genes of 638R compared to 638R-GA1Δ*acrR*_GA1_ from feces of mice monocolonized with each strain. **(D)** Competitive colonization assay in gnotobiotic mice competing 638R versus 638R-GA1Δ*tssBC*_GA1_ in three female (F) and three male (M) mice. The percentage of the two strains in the initial inoculum and then in the feces at day 14 and day 28 post-gavage is shown.

Previous studies demonstrated potent antagonism by the GA3 T6SS of 638R and other *B. fragilis* strains *in vivo* (*10–12*), yet the *in vivo* competitive colonization assay between 638R and 638R-GA1 (Fig. 3A) revealed that 638R-GA1 outcompetes the ancestral 638R strain in the gnotobiotic mouse gut (Fig. 3A). Based on our finding that 638R-GA1 does not transcribe the GA3 T6SS genes *in vivo*, this strain would also not be able to protect itself from GA3-mediated antagonism. This result raises the possibility that the GA1 T6SS may be antagonistic *in vivo* and may be effective at competing against the GA3 weapon, especially when delivered from the 638R isogenic strain. This is consistent with the finding that the GA1 *hcp* gene and other GA1 T6SS genes are substantially more highly expressed *in vivo* than *in vitro* (datasets S1, S2). To test if GA1 antagonism may explain the 638R-GA1 competitive colonization advantage, we used the 638R-GA1 mutant in which the GA1 *tssBC*, encoding both GA1 sheath proteins, were deleted, which abrogates GA1 T6SS firing (Fig. 2E). Colonization assays competing this mutant, (638R-GA1Δ*tssBC*_GA1_), against WT 638R showed that 638R-GA1Δ*tssBC*_GA1_ is outcompeted by 638R (Fig. 4D). These data, combined with those of the 638R x 638R-GA1 competitive colonization data show that an active GA1 T6SS in a GA3-containing *B. fragilis* strains allows it to outcompete the isogenic ancestral strain *in vivo*.

This study shows that transfer of a GA1 ICE into GA3-containing *B. fragilis* strains inactivates GA3 antagonism at two levels. In broth cultures, the GA3 locus is transcribed, but not able to fire, whereas in the mammalian gut, the GA1 ICE inhibits transcription of the GA3 T6SS genes involving the AcrR_GA1_ transcriptional repressor. AcrR-family transcriptional repressors bind specific ligands resulting in their removal from their DNA targets, relieving repression. Under conditions when such de-repression occurs, the second level of antagonistic repression, i.e., the inhibition of firing, which is not AcrR-dependent, ensures the GA3 system remains inactive.

The extensive transfer of the GA1 ICE among diverse GA3-sensitive Bacteroidales species (*9*) increases opportunities for its transfer into GA3-containing *B. fragilis* strains. Antagonism mediated by the newly acquired GA1 T6SS then allows the *B. fragilis* GA1-transconjugant to outcompete the progenitor *B. fragilis* strain, thereby changing a major Bacteroidales-specific antagonist to one that no longer antagonizes other GA1-containing species of the ecosystem. The ecological consequences and fitness outcomes of this DNA transfer to both the recipient and the Bacteroidales community are likely dependent on numerous factors including the composition of the community, how many of its Bacteroidales species contain the GA1 ICE, the diet of the host as relates to competition for mucin glycans, and potency of the GA1 T6SS. The dynamics of which weapon would win in a battle of *B. fragilis* strains is likely to be highly strain dependent. Some Bacteroidales strains contain mobile genetic elements containing acquired interbacterial defense systems (*16*), which contain genes encoding immunity proteins to GA1 and GA2 toxic effectors. As GA1 ICEs continue to spread in human gut microbiomes, interbacterial defense islands that counteract toxic effectors of the GA1 T6SS may prevent the GA3 T6SS from becoming obsolete.

To our knowledge, this is the first demonstration of a MGE of Bacteroidales drastically altering a phenotype encoded by the recipient organism’s genome. As the catalog of mobile genetic elements of the gut microbiota and analyses of their phenotypes increases (*15a*,*16, 26-31*), it is important to consider that the effects these elements have on the native genome may be as ecologically relevant as the phenotypes encoded by the MGE themselves.

## Acknowledgements

We are grateful to R. Smith, C. Metcalfe, C. Woodson, and V. Burgo of the DFI Microbiome Metagenomics Facility for RNA sequencing and whole genome sequencing. We thank K. Hutt, K. Kolar, K. Campbell, J. Kasper and B. Theriault of the UChicago GRAF center for assistance with gnotobiotic mouse experimentation and S. Light for helpful discussions. We thank H. Chu for providing strain YCH46. A few strains used in this study were obtained through BEI Resources, NIAID, NIH as part of the Human Microbiome Project.

## Data and materials availability

All materials created in this study are available from the corresponding author. Constructs, strains and mutants are available with a materials transfer agreement (MTA). The genomes sequenced as part of this study and RNASeq data have been deposited with NCBI under BioProject ID PRJNA982086.

## Supplementary Materials

Datasets S1 to S2

References 32-44 are cited only in methods and supplemental materials

## Materials and Methods

All primers used in this study are listed in Table S3.

### Strains and growth conditions

All bacterial strains used in this study are listed in Table S4. Bacteroidales strains were grown in basal liquid medium (*32*) or on BHIS plates (*33*) under anaerobic conditions. Antibiotics used for selection or induction include erythromycin (10 µg/mL), gentamicin (200 µg/mL), tetracycline (4.5 µg/mL), anhydrotetracycline (75 ng/mL), cefoxitin (10 µg/mL), carbenicillin (100 µg/mL). The *B. fragilis* 638R strain used throughout this study is erythromycin resistant as described below.

### SDS-PAGE and western blot analyses

Cell lysates and supernatants analyzed in western immunoblots were from overnight cultures grown in basal medium. For cellular lysates, cells from 5 µl of culture were loaded into each well and 10 µl of supernatants were loaded. These were prepared in 4x NuPAGE LDS Sample Buffer (Invitrogen, Waltham, MA) and samples were reduced with Bolt sample reducing agent (Invitrogen). Samples were boiled for 5 minutes then run on 12% Bolt SDS-PAGE Bis/Tris gels (Invitrogen). Contents of SDS-PAGE gels were transferred to polyvinylidene difluoride (PVDF) membranes and then blocked with TBST (100 mM Tris, 0.9% NaCl, 0.1% Tween 20, pH 7.5) containing 4% skim milk for 15 min.

Mouse fecal samples were prepared for western immunoblot as follows: the fecal pellets from each mouse were diluted 1:10 w/v in basal medium and physically disrupted to liberate the bacteria. The fibrous material was sedimented by gravity for 5 min and 200 µl of the non-sedimented liquid was centrifuged to collect the bacteria and resuspended in 200 µl 1X NuPAGE LDS Sample Buffer with Bolt Sample reducing agent and boiled for 5 min. 10µl of this material was loaded on 12% Bolt SDS-PAGE Bis/Tris gels and processed for western immunoblot as described above.

Antiserum to Hcp_GA3_ and BSAP-1 were previously described (*4, 10*). Antiserum to Hcp_GA1_ was generated in rabbits at Lampire Biological Laboratories using the Expressline protocol using purified His-Hcp_GA1_ as immunogen. The use of rabbits for antiserum generation was approved by the Institutional Animal Care and Use Committee (IACUC), University of Chicago and complies with all relevant ethical regulations for animal testing and research. The Hcp_GA1_ antiserum was reactive to the Hcp_GA3_ protein so cross-reactivity was removed by adsorption (fig. S8). Briefly, the antiserum to Hcp_GA1_ was diluted 1:10 in TBS and applied to an Ni-NTA agarose column (Invitrogen) bound with His-tagged Hcp_GA3_ and inverted end-over end at room temperature for 30 minutes. The flow through portion from the column (adsorbed antiserum) (fig. S8) was used as Hcp_GA1_ antiserum for all analyses. Secondary antibody (α-rabbit IgG-alkaline phosphatase) was used at a 1:2000 dilution. The membranes were developed with BCIP/NBT (5-bromo-4-chloro-3-indolylphosphate/ Nitro Blue Tetrazolium) (Seracare Life Sciences, Milford, MA).

### GA1 transfer assays – antagonism assays

*tetQ* was added to the *B. finegoldii* CL09T03C10 (BfineCL09) (*1*) GA1 ICE between genes QR305_02979 and QR305_02980 to allow for selection of transconjugants. For this, *tetQ* was amplified from *Bacteroides caccae* CL03T12C61 (*1*) and cloned with the left and right flanks of the insertion site, into the PstI site of pLGB13 (*33*). Transformants were mated with BfineCL09, and cointegrants were selected with gentamicin and erythromycin. Cointegrants were passaged in basal without antibiotics for several hours and plated on BHIS aTC plates. Mutants were selected among the double cross-outs by PCR and by tetracycline resistance.

The *ermG* gene was added to the *B. fragilis* 9343, *B. fragilis* 638R, and *B. fragilis* 638R ΔV1ΔV2 genomes by the same method used to insert this gene into *B. fragilis* 638RΔT6 (*28*).

GA1 conjugal transfer and antagonism assays were quantified from the same co-cultures. BfineCL09 GA1-*tetQ* served as the donor/targeted strain and *B. fragilis* 638R, *B. fragilis* 638R ΔT6 (GA3), or *B. fragilis* 9343, all containing the *ermG* chromosomal insertion, as the recipient/antagonistic strain. Strains were grown in basal medium to an OD_600_ of 0.7 and mixed at ratios of (1:10, 1:5, 1:1, 5:1 and 10:1, donor:recipient). 10 µL of the mixed strains was plated on a BHIS plate for 19 h. The bacterial growth was resuspended in 1 mL basal and 100 µL was plated on BHIS with tetracycline and erythromycin to quantify transconjugants, and 10 µl of 10-fold serial dilutions (10^-4^-10^-8^) were plated on BHIS containing erythromycin to calculate the number of recipients. Transfer frequency is calculated as the number of transconjugants/number of recipients. For quantification of antagonism, dilutions were plated on tetracycline to select for BfineCL09 and the amount surviving co-culture with 638R versus 638RΔT6 allows for quantification of antagonism. fig. S3 shows the plates from one of two biological replicates.

### Genome sequencing and assembly

The genomes sequenced as part of this study have been deposited with NCBI under BioProject ID PRJNA982086.

Long-read sequencing of *B. finegoldii* CL09T03C10 (BioSample ID SAMN35685248), *B. fragilis* 638R-GA1(6) (SAMN35685252), *B. fragilis* 078320-1 (SAMN35685249) and *B. fragilis* S36-L11 (SAMN35685254) was performed by SNPsaurus (Institute of Molecular Biology, Eugene, OR) using the PacBio Seqel II HiFi platform and were assembled using Flye (version 2.9) (*34*).

Oxford NanoPore and Illumina MiSeq sequencing for *B. fragilis* 2_1_56FAA (SAMN35685251), *B. fragilis* 1284 (SAMN35685250), and *B. fragilis* Korea-419 (SAMN35685253) were performed by The Duchossois Family Institute Microbiome Metagenomics Facility (DFI-MMF, University of Chicago, Chicago, IL). Hybrid assembly of these genomes was achieved using Unicycler (v0.4.8) (*35*).

For all genomes, gene calls and annotation were performed using Prodigal 2.6.3 (*36*) and Prokka 1.14.6 (*37*), respectively.

### RNA recovery and sequencing

For RNA sequencing of broth grown strains, bacteria were grown in triplicate in basal medium to OD_600_ of 0.9 and the bacteria were added to microfuge tubes in the anaerobe chamber prior to centrifugation so as not to expose to oxygen. The resulting pellets were flash-frozen and sent to Novogene (Sacramento, CA). RNA was extracted, rRNA depleted, libraries were constructed, and RNA sequencing was performed (2 x 150 bp paired end) using the Illumina NovaSeq 6000 platform.

For RNA sequencing from mouse feces, three male mice for each inoculum (representing biological triplicates) were gavaged with the following *B. fragilis* strains, all with the chromosomal *ermG* insertion and the *tetQ* deleted from the GA1 ICE for the last two strains: 638R, 638R-GA1-6 or *B. fragilis* 638R-GA1Δ*acrR*_GA1_. Fecal pellets were collected from each mouse on day 7 post-gavage and immediately placed on dry ice and frozen at -80°C until processed by the DFI-MMF. Briefly, fecal samples were first subjected to bead-beater disruption using a TissueLyzer II bead mill (Qiagen). Nucleic acid was recovered using the Maxwell RSC instrument (Promega). After DNAse treatment and elution, the samples were quantified by measurement on a Qubit (Life Technologies) and integrity was assessed by use of a Tapestation unit (Agilent Technologies). Ribosomal RNA was depleted from all samples with use of the NEBNext rRNA depletion kit (New England Biolabs). The samples were fragmented, libraries were prepared using the Ultra Directional RNA library prep kit for Illumina (New England Biolabs), and normalized libraries of biological triplicates of all samples were sequenced on Illumina’s NextSeq 1000 platform at 2 x 100 bp read length.

### RNASeq analyses

After adapter and quality trimming of all reads using utilities included in the BBMap package of bioinformatics tools (v. 38.90), the reads were mapped to the *B. fragilis* 638R-GA1 (6) genome assembly using the Bowtie 2 short-read aligner (v. 2.4.2) (*38*). The Bowtie 2 output was converted to sorted and indexed BAM files using SAMtools (v. 1.11) (*39*), and BEDtools (v. 2.30.0) (*40*) was used to compare these to a General Feature Format (GFF) file containing the intervals of protein-coding domains from *B. fragilis* 638R-GA1 (6). The read mapping results were evaluated for differential gene expression using both DESeq2 (v. 1.30.0) (*41*) and edgeR (v. 3.32.1) (*42*). A gene was considered differentially expressed if both statistical packages agreed that the absolute value of its fold change (FC) in expression level under experimental conditions differed from the control conditions by ≥2 and if the adjusted *p*-value (padj for DESeq2 and FDR for edgeR) was ≤ 0.05. edgeR calculations were relied on exclusively for determination of differential expression in cases where DESeq2 returned NA due to read count outlier detection. Volcano plots were created in Prism, version 9.5.1 (GraphPad Software, San Diego, CA). RNASeq data were deposited with NCBI under BioProject ID PRJNA982086.

### Mutant construction, complementation and gene addition

DNA regions for all constructs were PCR amplified using Phusion polymerase (NEB) with primers listed in Table S4. NEBuilder (NEB) was used for all cloning and assembly of DNA segments into vectors. Whole plasmid sequencing was performed for all recombinant plasmids by Plasmidsaurus (Eugene, OR) or Primordium (Monrovia, CA).

#### Insertion of ermG into B. fragilis 638R and B. fragilis 9343

A construct that was previously created to insert *ermG* into the chromosome of 638RΔT6 between genes Bf638R_0867 and Bf638R_0868 (*28*) was used for scarless double crossover insertion of the *ermG* into WT 638R and 9343 to allow for selection of transconjugants receiving the GA1 ICE or for selection in antagonism assay, or into 638RΔV1ΔV2 (*10*)(deleted for the two variable regions of the GA3 T6SS) for selection in antagonism assays (below).

#### Insertion of tetQ into the BfineCL09 GA1 ICE

*tetQ* was added into the BfineCL09 GA1 ICE between genes QR305_02979 and QR305_02980. *tetQ* was amplified from *B. caccae* CL03T12C61 (HMPREF1061_01202) and cloned with the left and right flanking regions into PstI digested pLGB13 (*33*) and then then transformed into *E. coli* S17 λ pir and plated on LB plates with carbenicillin. The correct sequence-confirmed plasmid was conjugally transferred from *E. coli* to BfineCL09 and cointegrates were selected on BHIS containing erythromycin and gentamicin. A cointegrate was passaged in basal medium lacking antibiotics for several hours and plated on BHIS with anhydrotetracycline to select for double crossout recombinants. Mutants were confirmed by PCR and acquisition of tetracycline resistance.

#### Construction of pMLS36 vector for mutant construction using cefoxitin resistance

Vector pMLS36 was constructed by replacing the *ermG* of pLGB36 (*43*) with *cfxA* (INE91_01220) of *Phocaeicola vulgatus* CL11T00C01 using the primers listed in Table S4.

#### Construction of deletion mutants

All GA1 ICE deletion mutants in 638R-GA1 were constructed by cloning the flanking segments of the region to be deleted into the BamHI site of pMLS36. Plasmids were conjugally transferred from *E. coli* to 638R-GA1 and co-integrates were selected on BHIS containing cefoxitin and gentamicin. Cointegrates were passaged in basal media without antibiotics for several hours and plated on BHIS anhydrotetracycline to select for double crossout recombinants. Mutants were identified by PCR. The deletion of *acrR*_GA1_ from 9343-GA1 was created using the same construct for deletion of this gene from 638R-GA1. Deletion of *acrR*_GA1_ (CQW33_03258) from *B. fragilis* 1284 was made by cloning the flanking regions into BamHI digested pLGB36. Deletion of Bf638R_1555-1552 from 638R*ermG* was made by cloning the flanking regions into pMLS36.

#### Construction of His-Hcp_GA1_

An N-terminal His-tagged fusion of Hcp_GA1_ was constructed by cloning QR305_02952 into the NdeI–BamHI sites of pET16b (Novagen), and the plasmid was transformed into *E*. *coli* BL21/DE3. Following induction, His-Hcp_GA1_ was purified using the ProBond Purification System (Life Technologies) and the eluted protein was dialyzed against PBS and used as immunogen for antiserum generation.

### Antagonism assays

All antagonism assays used a 10:1 ratio of antagonist to target cell except for the combined ICE transfer-antagonism assays shown in fig. S3, where different ratios were used. All cultures were grown to an OD_600_ of ∼0.7, mixed, and 10 µl was dotted to a BHIS plate and incubated 18-20 hours under anaerobic conditions. The bacterial growth was resuspended in basal medium and 10 µl l of 10-fold serial dilutions were plated. For assays where 638R, 638R-GA1(5), 638R-GA1(6), 638R-GA1Δ3, 638R-GA1Δ4, 9343, 9343-GA1 (2b), and 9343-GA1 (3a) were the antagonist, *B*. *thetaiotaomicon* VPI-5482 served as the target strain and the serial dilutions were plated on BHIS with cefoxitin, which selects for *B. thetaiotaomicron* VPI-5482. The assay shown in Fig. 1F to test for antagonism of the 638R-GA1 strains by progenitor strain 638R shows the killing of an isogenic strain, 638RΔV1ΔV2 (deleted for both variable regions encoding the effectors and immunity proteins of the GA3 locus so is not able to protect itself), by the 638R WT strain, but not by the 638R ΔT6 strain that is unable to fire the GA3 T6SS (*10*).

### Gnotobiotic mouse experiments

All mouse experiments were approved by the Institutional Animal Care and Use Committee (IACUC), University of Chicago. All mice were bred at the Gnotibiotic Research Animal Facility (GRAF) at the University of Chicago. After inoculation, mice were housed in a cage rack system to maintain gnotobiotic status. All mice were germ-free C57BL/6J mice age 6-10 weeks at the time of gavage.

For monocolonization assays, male mice were gavaged with 200 µl of log-phase grown bacteria in basal medium. Feces were collected and analyzed seven days post-gavage.

For competitive colonization experiments, *tetQ* was removed from the GA1 ICE of all strains where the ICE was transferred *in vitro*, to restore the GA1 ICE to its native state. The two competing strains were grown to an equal OD_600_ and 200 µl of a 1:1 volume mixture was gavaged into three female and three male mice housed separately by sex. The initial inoculum ratio was experimentally quantified. Feces were collected under sterile conditions over time and the ratio of strains was quantified by multiplex PCR.

---

**Fig. S1.**
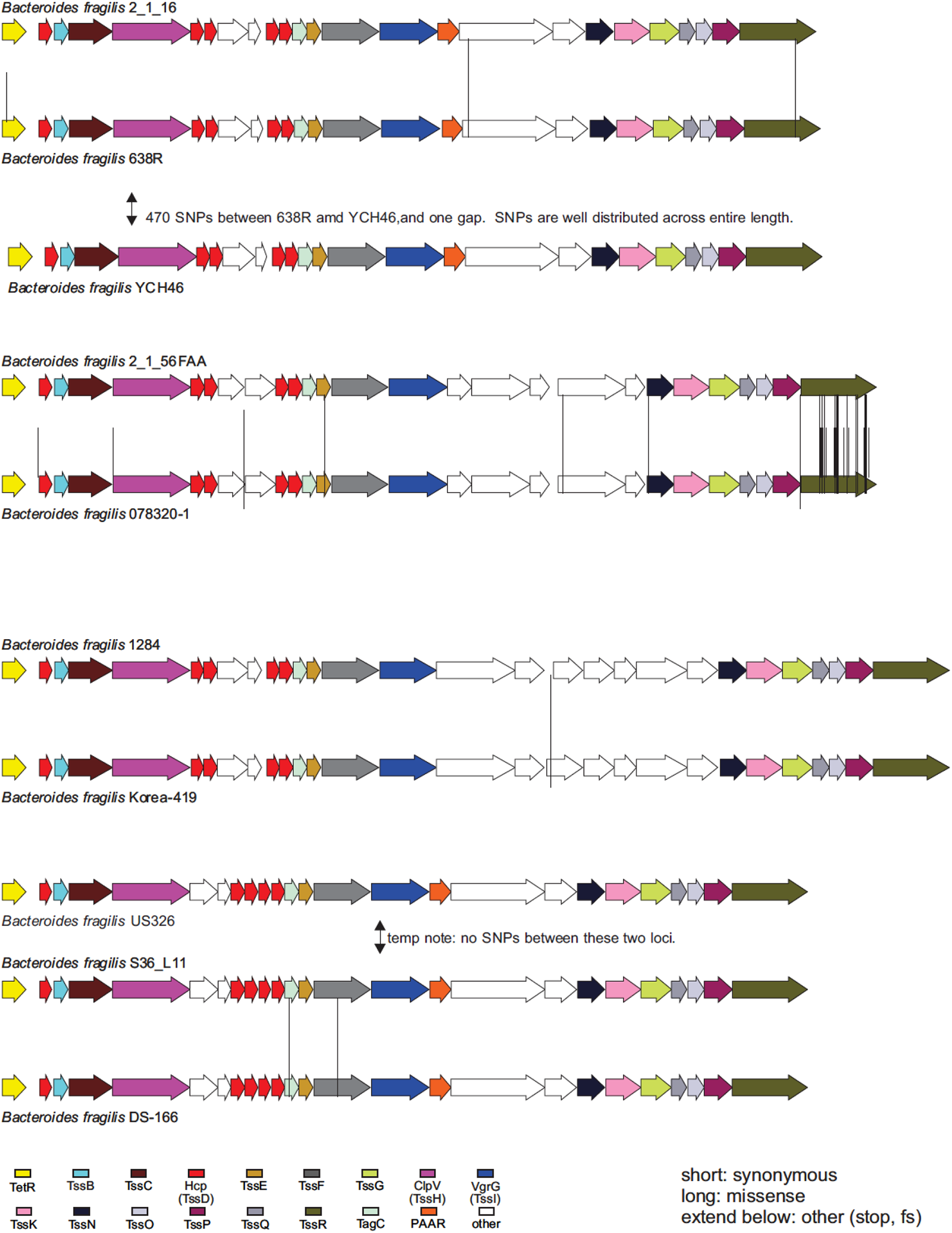
Schematic of the nucleotide differences in the GA3 loci of *B. fragilis* strains that have the same V1 and V2 regions. Short vertical lines represent synonymous differences, those that extend the full length between the regions are sites of non-synonymous differences, and those that extend below the bottom gene map are differences conferring a stop codon or frame shift. None of these regions have any obvious defects in essential T6SS genes.

**Fig. S2.**
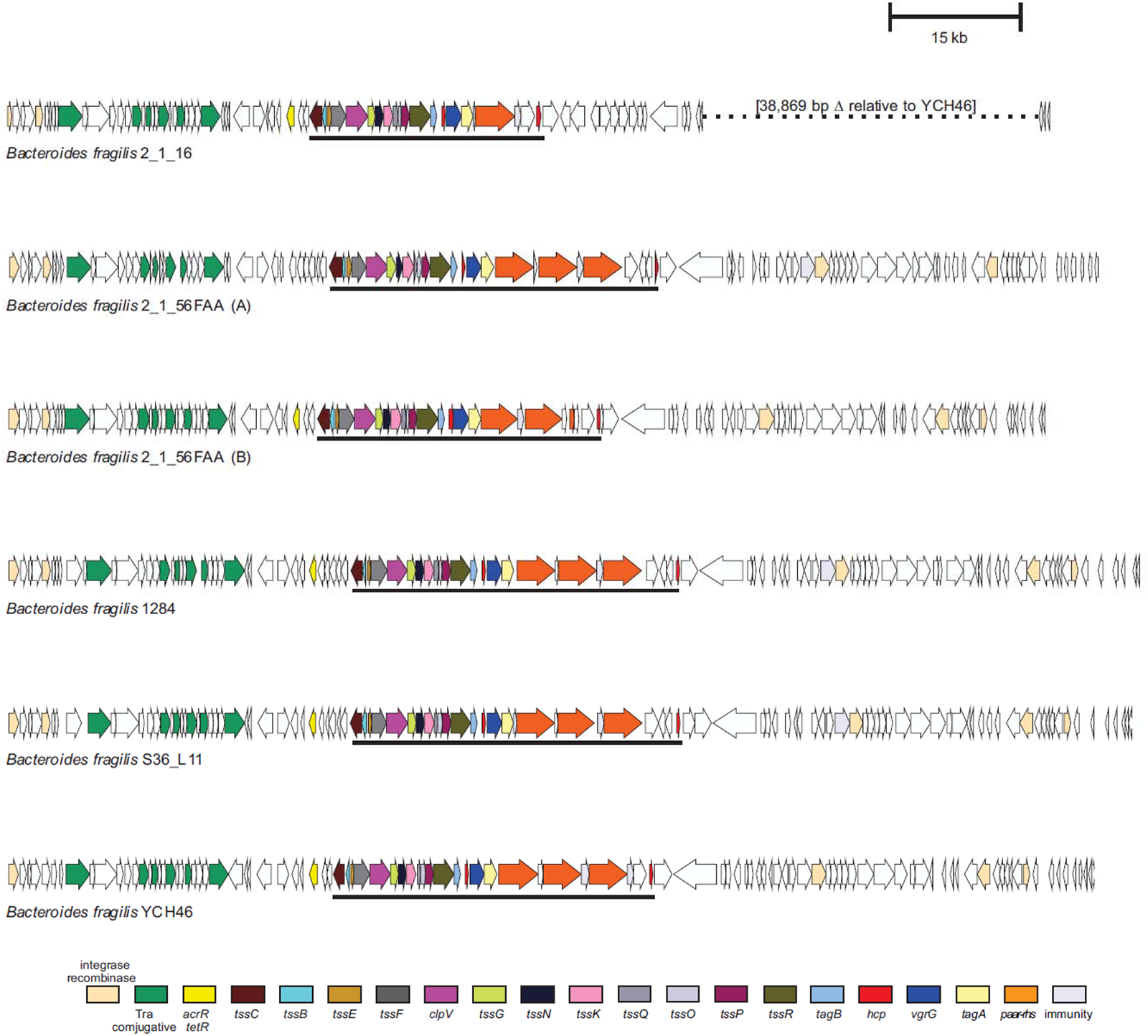
Genetic maps of the GA1 ICE contained in the six *B. fragilis* strains shown in panels A and B. Annotations of the genes of the regions are shown in the color-coding below. The GA1 ICE of *B. fragilis* 2_1_16 is missing a large region at the right side of the GA1 ICE.

**Fig. S3.**
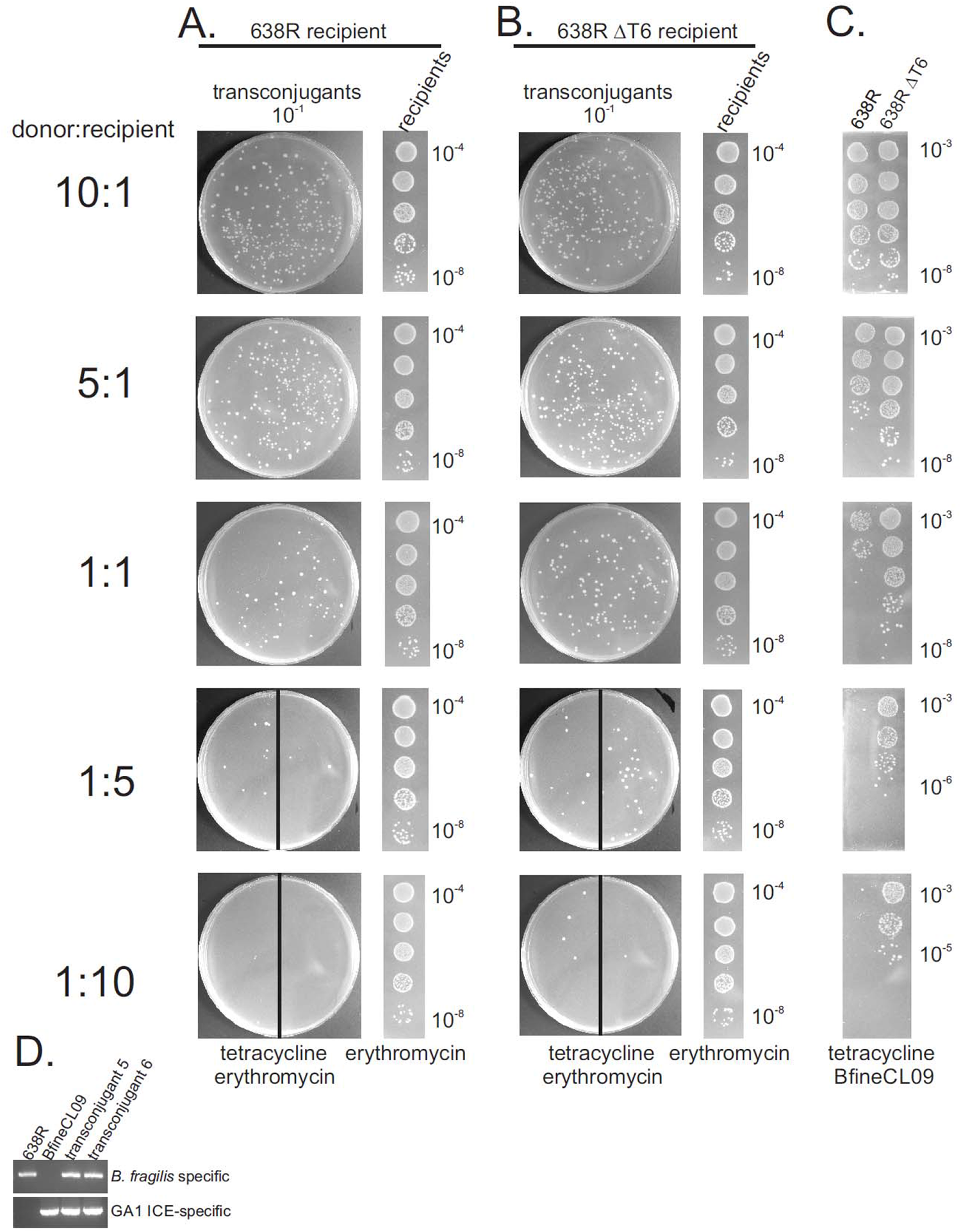
GA1 ICE transfer from *B. finegoldii* CL09T03C (BfineCL09) to 638R or 638RΔT6. **(A), (B)** Transfer assay plates showing number of transconjugants (tetracycline and erythromycin resistant) on the 10^1^ dilution plate and dilutions of recipients 638R (column A) or 638RΔT6 (column B) (erythromycin resistant) at five different ratios of donor:recipient. For 1:5 and 1:10 ratios, biological replcates of transconjugants were plated on half a plate due to low abundance **(C)** Antagonism assay showing survival of BfineCL09 from the same co-culture as the ICE transfer assay.**(D)** PCR analysis showing that transconjugants are the 638R background containing the GA1 ICE.

**Fig. S4.**
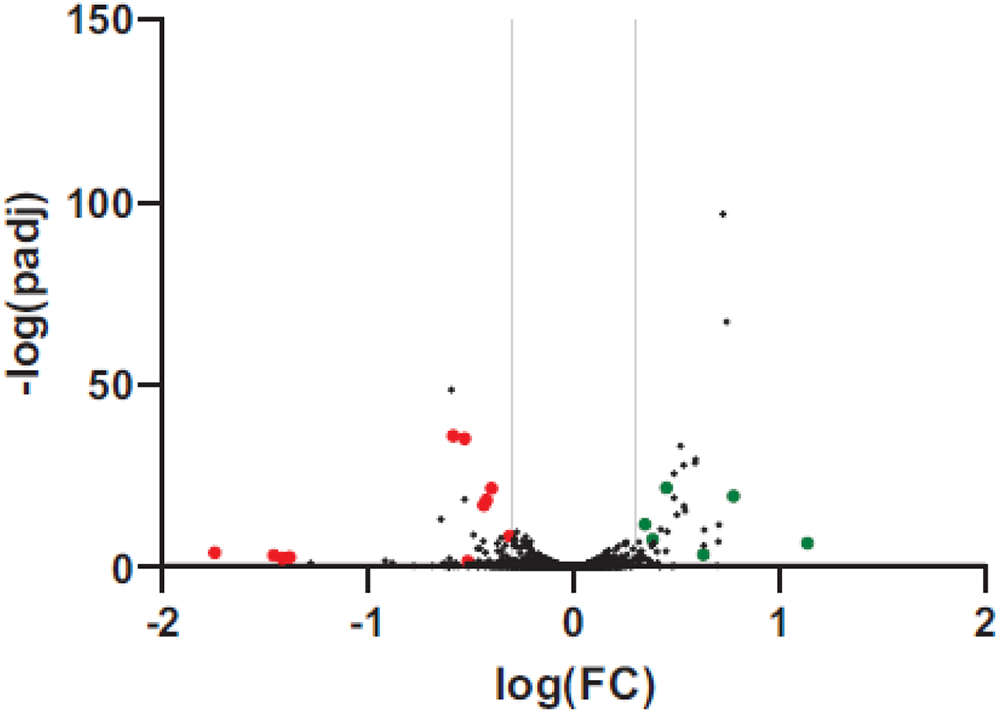
Volcano plot showing the· differentially expressed genes of 63ARGA1 transconjugant 5 compared to 638R. Red and green dots show genes that were also similarly differentially expressed in 638RGA1 transconjugant 6. Very few genes are differentially expressed under these in vitro conditions. Full RNAseq data provided In Table S1.

**Fig. S5.**
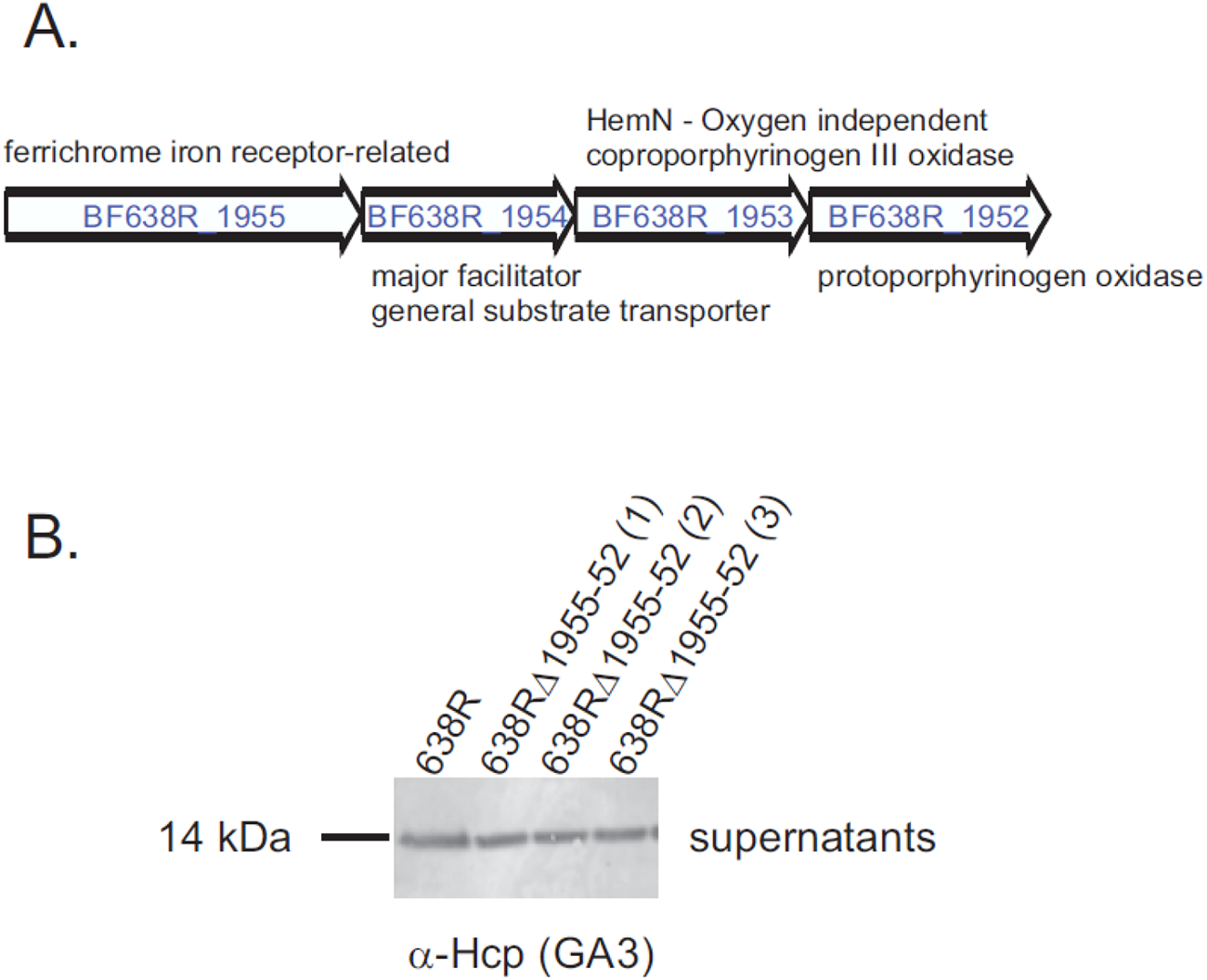
An operan of four genes all significantly downregulated in 638R-GA1 compared to WT 638R grown in basal medium. **(A)** genetic organization showing the predicted functions of the proteins encoded by each gene. **(B)** Western immunoblot showing the secretion of the GA3 Hep from 638R deleted for these four genes. Three isogenic mutants are shown.

**Fig. S6.**
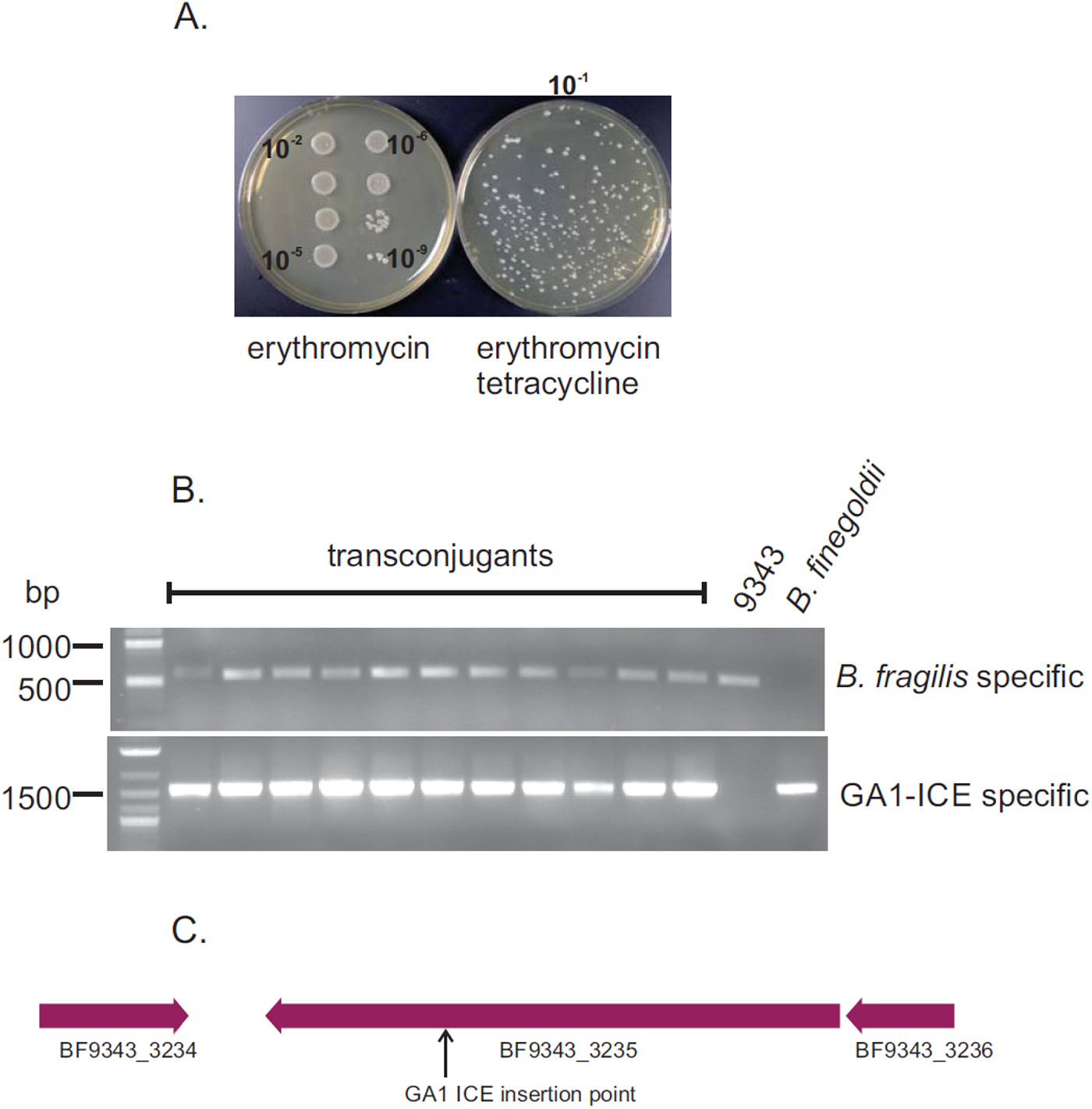
Conjugal transfer of the GA1 ICE from *B. finegoldii* CL09T03C10 to *B. fragilis* 9343. **(A)** Following co-culture at a 10:1 dononrecipient ratio for 19 hours, the erythromycln/tetracycline plate shows the number of transconjugants and the erythromycin plates shows the number of 9343 recipients. **(B)** Eleven random colonies from the erythromycin-tetracycline plate were restruck to a fresh plate and after regrowth were PCR screened with a primer set specific to *B. fragilis* and a second set specific to the GA1-ICE. The agarose gel of the PCR products shows that all are 9343-GA1 transconjugants. **(C)** Insertion site of the GA1 ICE into the 9343 chromosome in 9343-GA1-2b. The ICE inserted Into the last gene of a putative fimbrial operon.

**Fig. S7.**
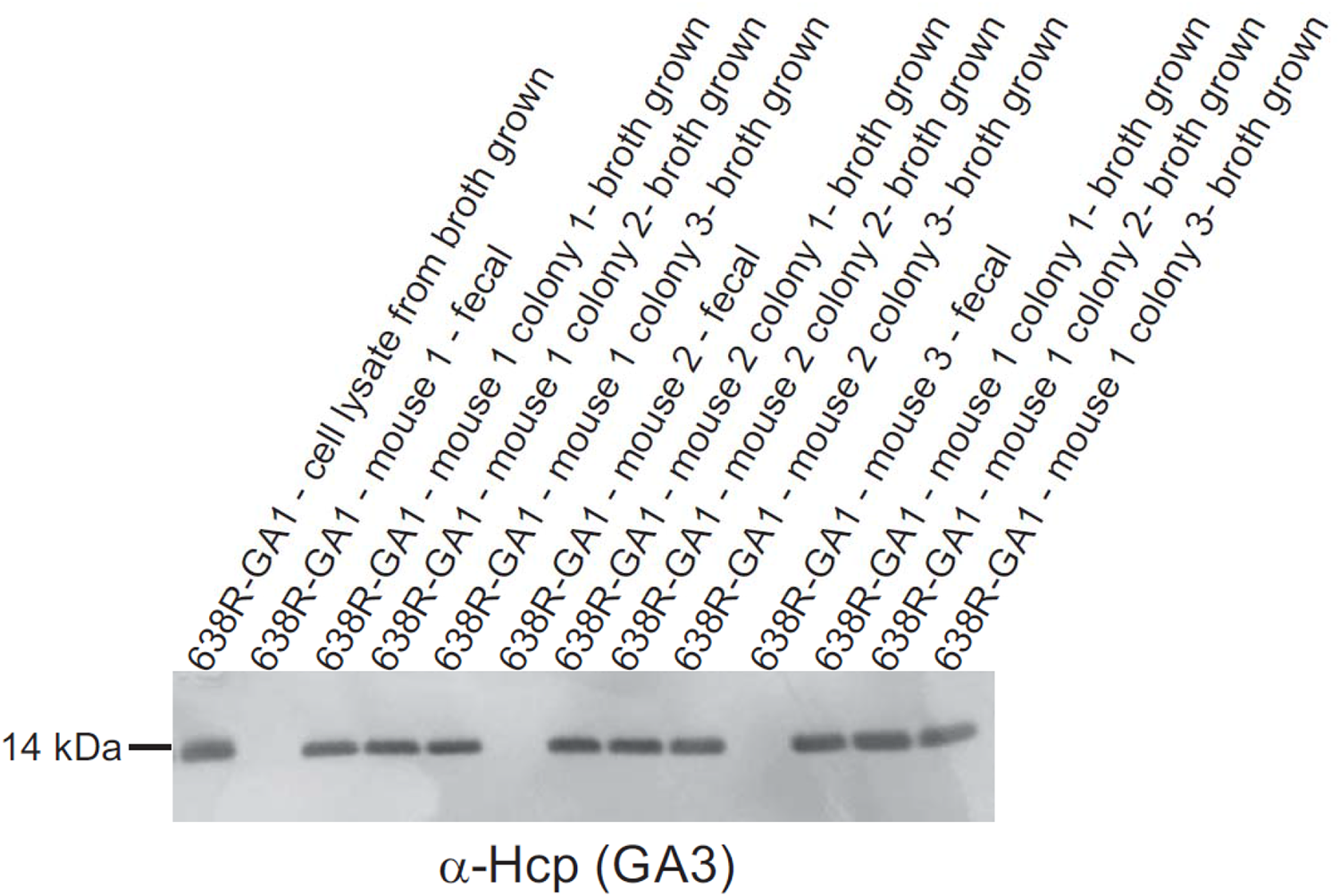
Analysis of GA3 Hep production from bacteria of monocolonized mice that were regrown in broth. Feces of mice monocolonized with 638R-GA1 were diluted and plated on BHIS plates and three resulting colonies from each mouse were passaged overnight in basal medium and the resulting cell lysate was analyzed by western blot with antiserum to GA3 Hep.

**Fig. S8.**
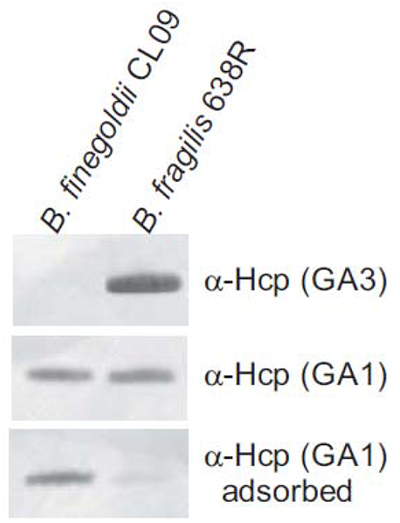
Reactivity of GA1 Hep antiserum before and after adsorption with GA3 Hep. Western immunoblot blot analysis of anti-GA3 showing it does not cross-react with GA1 Hep of *B. finegoldii* CL09T03C10, which was grown in defined minimal medium with *N*-acetyl glucosamine as the sole carbon source for expression of the GA1 Hep. Adsorption substantially removed cross-reactivity with the GA3 Hep.

**Table S1.**
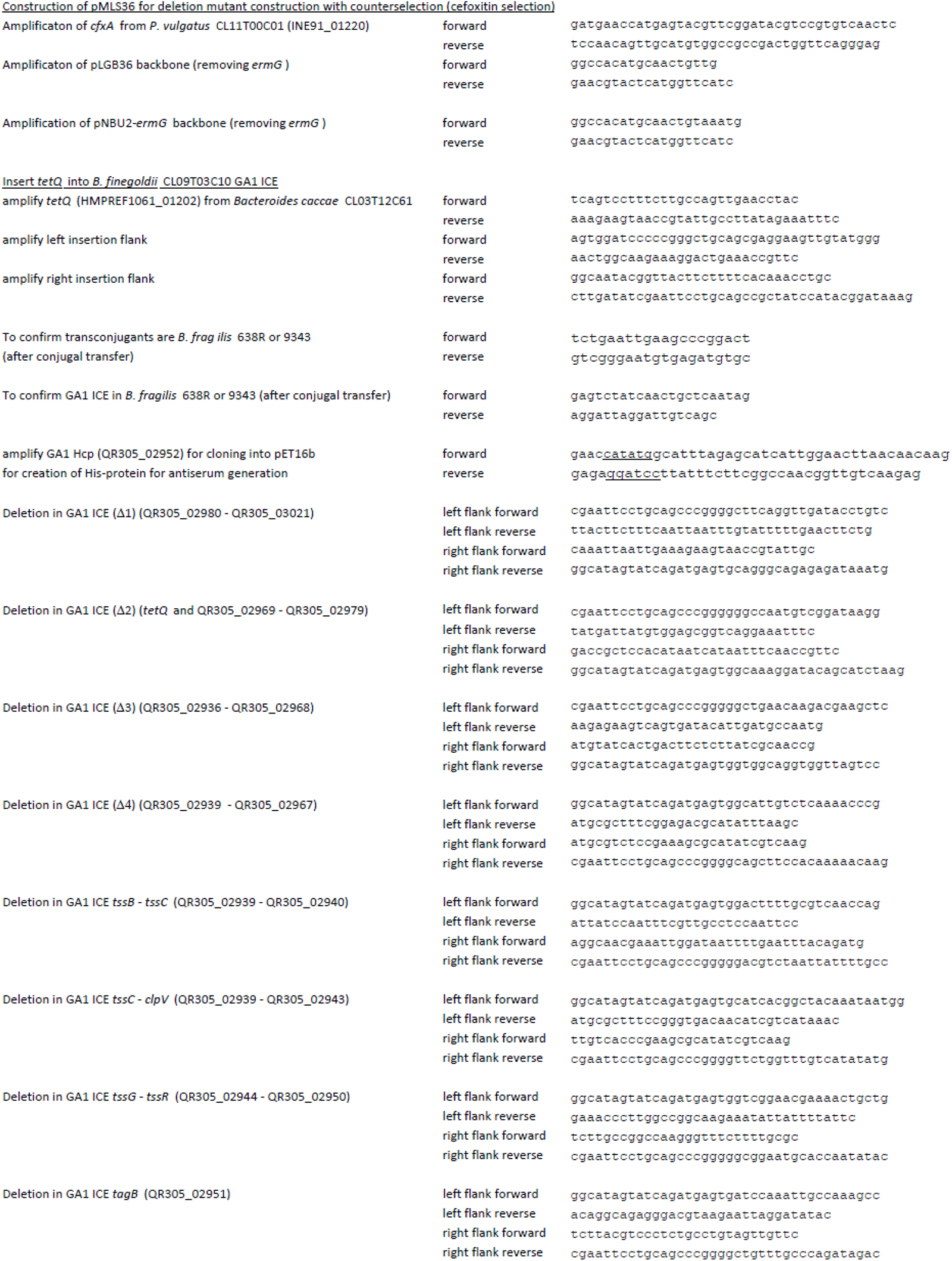
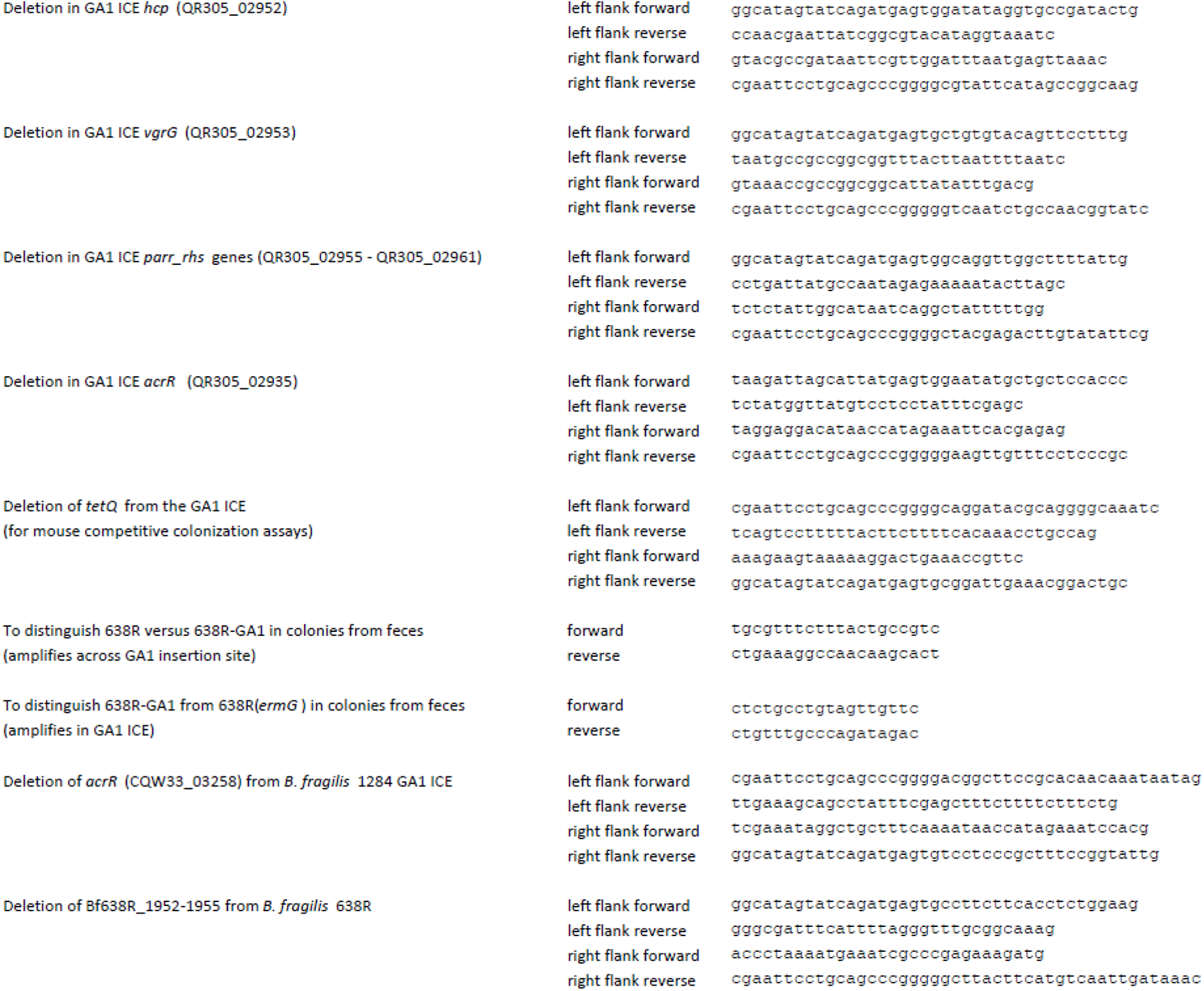
Primers used in this study.

**Table S2.** Strains used in this study.

| Table S2. Strains used in this study | source/reference |
| --- | --- |
| *Bacteroides fragilis* 638R | lab stock |
| *Bacteroides fragilis* 2_1_16 | BEI |
| *Bacteroides fragilis* YCH46 | ref 44 |
| *Bacteroides fragilis* CL07T12C05 | ref 1 |
| *Bacteroides fragilis* Korea 419 | lab stock |
| *Bacteroides fragilis* 1284 | lab stock |
| *Bacteroides fragilis* NCTC9343 | lab stock |
| *Bacteroides fragilis* 078320-1 | lab stock |
| *Bacteroides fragilis* 2_1_56FAA | BEI |
| *Bacteroides fragilis* J38-1 | lab stock |
| *Bacteroides fragilis* I-1345 | C. Sears |
| *Bacteroides fragilis* DS166 | BEI |
| *Bacteroides fragilis* S36_L11 | BEI |
| *Bacteroides thetaiotaomicron* VPI 8452 | lab stock |
| *Bacteroides caccae* CL03T12C61 | ref 1 |
| *Phocaeicola vulgatus* CL11T00C01 | ref 9 |
| *B. fragilis* 638R ΩermG | this study |
| *B. fragilis* 638R ΔT6 (ΔT6 is deletion of BF638R_1991-1993) | ref 10 |
| *B. fragilis* 638R ΔT6 ΩermG | ref 28 |
| *B. fragilis* 638R ΔV1ΔV2 (deletion of two effector/immunity regions of GA3 T6 locus) | ref 10 |
| *B. fragilis* 638R ΔV1ΔV2 ΩermG | this study |
| *Bacteroides finegoldii* CL09T03C10 | ref 1 |
| *B. finegoldii* CL09T03C10 GA1-ΩtetQ | this study |
| *B. fragilis* 638R ΩermG (GA1<sub>Bfine</sub> - ΩtetQ) (5) | this study |
| *B. fragilis* 638R ΩermG (GA1<sub>Bfine</sub> - ΩtetQ) (6) a.k.a. Bf638R-GA1 | this study |
| Bf638R-GA1 ΔtetQ (for mouse experiments) | this study |
| Bf638R-GA1 Δ1 (QR305_02980-QR305_03021) | this study |
| Bf638R-GA1 Δ2 (tetQ and QR305_02969 - QR305_02979) | this study |
| Bf638R-GA1 Δ3 (QR305_02936 - QR305_02968) | this study |
| Bf638R-GA1 Δ4 (QR305_02939 - QR305_02967) | this study |
| Bf638R-GA1 ΔtssB-tssC (QR305_02939 - QR305_02940) | this study |
| Bf638R-GA1 ΔtssB-tssC ΔtetQ (for mouse experiments) | this study |
| Bf638R GA1 ΔtssC-clpV (QR305_02939 - QR305_02943) | this study |
| Bf638R-GA1 ΔtssG-tssR (QR305_02944 - QR305_02950) | this study |
| Bf638R-GA1 ΔtagB (QR305_02951) | this study |
| Bf638R-GA1 Δhcp (QR305_02952) | this study |
| Bf638R-GA1 ΔvgrG (QR305_02953) | this study |
| Bf638R-GA1 Δrhs (QR305_02955 - QR305_02961) | this study |
| *B. fragilis* NCTC 9343 ΩermG | this study |
| *B. fragilis* NCTC 9343 ΩermG (GA1<sub>Bfine</sub> - ΩtetQ) (2) a.k.a. Bf9343-GA1 | this study |
| *B. fragilis* NCTC 9343 ΩermG (GA1<sub>Bfine</sub> - ΩtetQ) (3) | this study |
| BF9343-GA1 ΔacrR<sub>GA1</sub> | this study |
| BF638R-GA1 ΔacrR<sub>GA1</sub> (QR305_02935) | this study |
| *B. fragilis* 1284 ΔacrR<sub>GA1</sub> (CQW33_03258) | this study |
| *E. coli* S17 $\lambda$ pir | E. Martens |

